# GEQO biosensors for absolute analyte quantification in single cells

**DOI:** 10.1101/2025.05.05.652245

**Authors:** Sascha M. Kuhn, Elisa Nerli, Jifeng Liu, Sylvia Kaufmann, Pavel Barahtjan, Martin Buitrago-Arango, Eric R. Geertsma, Rita Mateus, Anne Grapin-Botton, André Nadler

## Abstract

Genetically encoded fluorescent biosensors are widely used to monitor small molecule and ion levels in living cells. Quantitative FRET and FLIM sensors and highly sensitive intensiometric sensors have been developed for many analytes. Notwithstanding notable advances over the last years, a universal high-performance sensor design for absolute quantification that does not require specialized equipment has yet to be developed. We here report the GEQO platform of quantitative biosensors featuring calcium, ATP, cAMP, and organelle-specific variants. We used GEQO sensors to follow calcium and cAMP transients in immortalised cells, human pancreatic progenitor cells, and zebrafish embryos. We show that GEQO-based absolute quantification allows to account for analyte buffering and retains information in time trace data lost during relative quantification. GEQO biosensors will enable quantitative analyte measurements across a wide range of imaging platforms, a key prerequisite for diagnostic applications and quantitative approaches in basic cell biology.

## Introduction

Genetically encoded fluorescent biosensors are essential for the investigation of molecular dynamics in living systems. Among the most popular biosensor classes are intensiometric reporters which translate changes in analyte concentration into changes in fluorescence intensity^1,2^. Their main advantages are high signal-to-noise ratios and wide dynamic ranges^3,4^. The range of analytes that can be detected is broad, including second messengers like calcium and cAMP, or metabolites like ATP and glucose^5–8^. Intensiometric measurements typically yield relative quantifications, as utilising the fluorescence readout for absolute analyte quantification requires knowledge of the sensor protein concentration^9,10^.

In order to routinely determine absolute analyte levels, biosensors have to allow for straightforward absolute quantification and results have to be comparable across imaging platforms. A number of biosensor types already allow for absolute analyte quantification. Ratiometric, typically Förster resonance electron transfer (FRET) sensors^11–13^, or fluorescence lifetime imaging microscopy (FLIM) sensors^14–17^ are the main examples. However, FRET sensors have a narrower dynamic range and poor signal-to-noise ratio compared to intensiometric sensors^18–21^, while FLIM sensors require specialised microscopes. Therefore, FLIM and FRET sensors are used much less frequently in cell biology compared to intensiometric biosensors. Ideally, genetically encoded fluorescent biosensors should exhibit the high signal-to-noise ratio and wide dynamic range of intensiometric biosensors while allowing for absolute quantification on all common imaging platforms.

Here, we introduce a family of **g**enetically **e**ncoded **q**uantitative **o**ptical indicators (GEQO) which are based on green sensor fluorescent proteins (FP) and an analyte-independent red long Stokes shift (LSS)^22^ reference FP. While the sensor FP provides high sensitivity in response to analyte concentration changes, the reference FP yields a direct read-out of the reporter’s expression level. Compared to previous attempts of realising this design, GEQOs feature minimal crosstalk between reporter and calibration channel^23,24^. Both channels can be independently calibrated, yielding a set of parameters for direct quantification of absolute analyte concentrations. The combination of a GFP sensor with a red LSS reference FP results in a ratiometric reporter that allows for single wavelength excitation. This feature simplifies microscope calibration and allows for straightforward applicability between imaging platforms ranging from plate readers to high-throughput screening microscopes (Fig. 1a). The modular sensor architecture enables converting established intensiometric sensors into GEQO sensors and organelle-specific variants. We demonstrated this by converting the cytosolic jGCaMP8f sensor into the ratiometric caGEQO and the plasma membrane targeted version caGEQO-PM. In addition, we generated the cAMP sensor campGEQO and the ATP sensor atpGEQO based on the intensiometric sensors cAMPinG1^6^ and iATPSnFR^8^.

**Figure 1.**
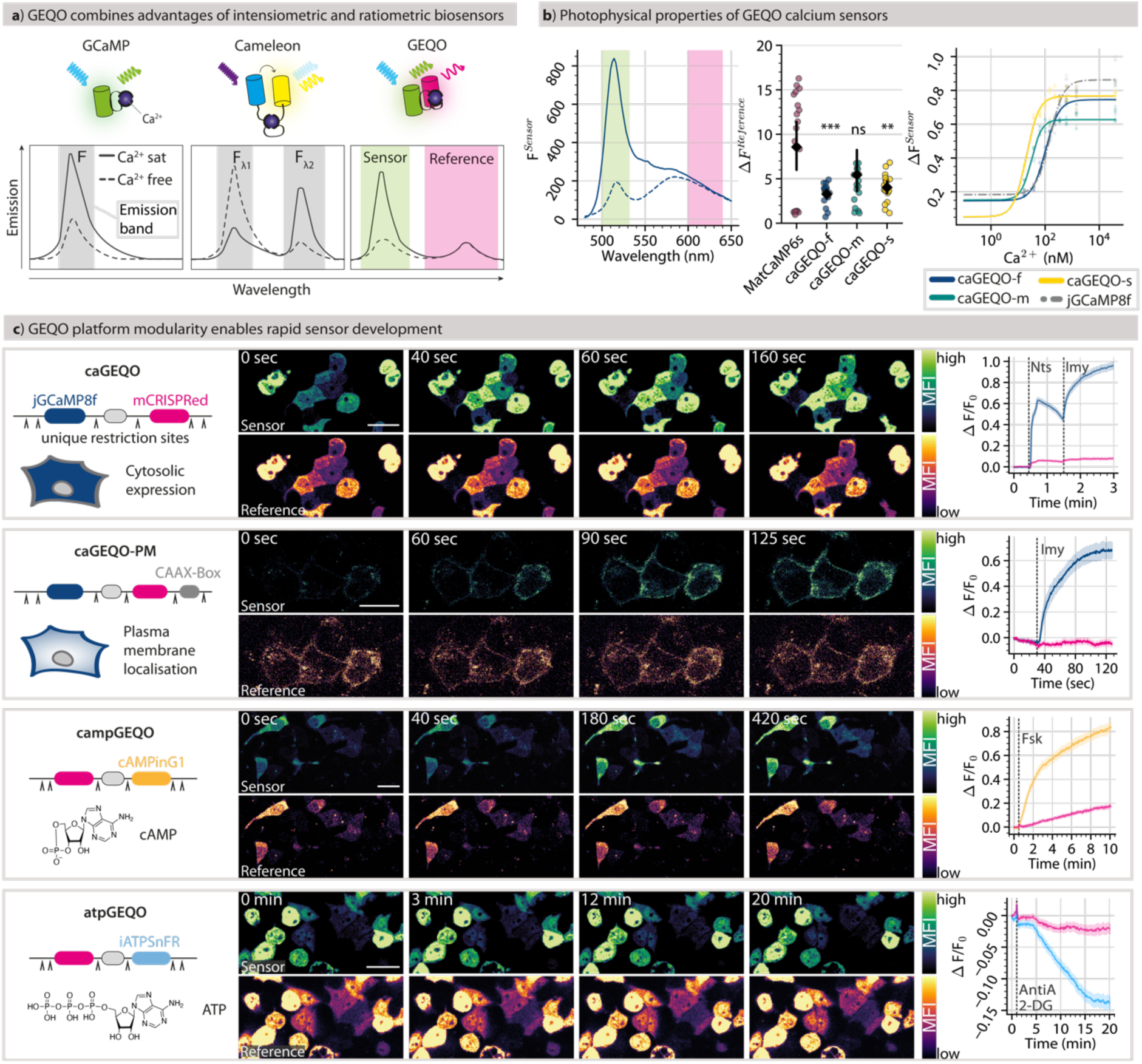
The GEQO sensor family is a versatile platform for measuring analyte concentrations inside cells. **a)** Schematic of different calcium sensor types. Intensiometric GCaMP-style sensors report calcium concentration changes in a single fluorescent emission band. In FRET-based ratiometric sensors, like Cameleon, two emission bands change in opposite directions. In GEQO sensors only one of two emission bands increases in response to calcium, the second emission band is insensitive to calcium. **b)** The emission spectrum of caGEQO2f *in-vitro* under calcium free and saturated (39 µM) conditions (left). Fluorescent bleed-through between sensor and reference emission bands *in-vitro* of MatryoshCaMP6s (MatCaMP6s, purple), caGEQO2f (blue), m (green), and s (yellow). Coloured dots represent individual measurements; mean and 95% confidence interval (bootstrapped) are shown in black; P-values of GEQO sensors in comparison to MatryoshCaMP are calculated using two-sided bootstrap hypothesis testing (***: p < 0.001, **: p < 0.01, ns: p > 0.05). **c)** Schematic of GEQO construct design (left). ^ indicate positions of unique restriction sites flanking the individual domains (coloured boxes). Middle panels show representative time lapse montages from HCT116 cells expressing the respective sensor (sensor channel in green hot, reference channel in inferno, scale bars: 25 µm). Right panels show relative quantification of the respective bleach corrected time series (solid line: mean, shaded area: standard error of the mean). 1^st^ row shows caGEQO-f (n = 154) comprising jGCaMP8f (dark blue) and mCRISPRed (magenta). Cells were treated with neurotensin (Nts, 100 nM after 30 s) and ionomycin (Imy, 5 µM after 90 s). 2^nd^ row shows the plasma membrane targeted version of caGEQO-f (n = 28), which incorporates a C-terminal CAAX-Box (dark grey). Cells were treated with Nts after 30 s. cAMP (3^rd^ row, n = 119) and ATP (4^th^ row, n = 272) sensitive GEQO versions were generated by exchanging jGCaMP8f for cAMPing1 (orange) and iATPSnFR (light blue), respectively. Stills and average time traces show the responses to forskolin (Fsk, 60 µM, after 30 s) and combined Antimycin A / 2-Desoxyglucose (AntiA 10 µM, 2DG 10 mM, after 30 s) treatments.

We used GEQO sensors to measure GPCR-induced calcium and cAMP responses in immortalised cells and established an analysis platform for classifying single cell responses. Using this pipeline, we developed a strategy for alleviating common artifacts of biosensor use such as analyte buffering, which results in artificially low baseline analyte levels and suppressed peak amplitudes of signaling transients. We quantified externally induced calcium transients in human pancreatic progenitor cells and followed spontaneous calcium oscillations in developing zebrafish embryos. Using these datasets, we show that both intensiometric and ratiometric data analysis led to information loss compared to absolute quantification, demonstrating the potential of the GEQO platform for in-depth and highly reliable determination of second messenger dynamics by fluorescence imaging

## Results

### GEQO design and photophysical properties

The design of the GEQO sensor platform combines the advantages of ratiometric and intensiometric biosensors. Excitation at a single wavelength (455 nm) results in a distinctive emission spectrum featuring two separate bands. A first emission band (sensor channel, 520 nm) is sensitive to analyte concentration, similar to intensiometric sensors. A second, analyte-insensitive emission band (reference channel, 610 nm) simultaneously enables ratiometric imaging and quantification of biosensor concentration, which is not straightforward for traditional FRET-based sensors^10,25,26^ (Fig. 1a). To obtain a sensor providing these properties, we constructed different fusion proteins containing the green intensiometric calcium indicators jGCaMP8f, m, or s^27^ in combination with the long Stokes shift fluorescent proteins (FPs) LSS-mKate2^28^ or mCRISPRed^29^. Sensors featuring the LSS-mKate FP exhibited only weak fluorescence in the red channel and were not further pursued (Extended Data 1a). We decided to pursue mCRISPRed-based sensors, as this FP offers a combination of relatively high brightness, quantum yield, relative pH insensitivity, and a comparably large Stokes shift. We found that the dynamic ranges of GEQO sensors were lowered in comparison to their parent sensors, presumably due to green fluorescence originating from the reference FP. The combination of jGCaMP8f with mCRISPRed gave the best-performing (Fig. 1b) ratiometric sensor, which we named caGEQO, for calcium sensitive GEQO. This sensor version showed the desired properties of minimal fluorescence intensity change in the reference channel (Figure 1b, c), high signal-to-noise ratio in cellular experiments (Extended Data 1b), and similar calcium binding affinity and emission spectra to its parent sensor jGCaMP8f (Figure 1b, Extended Data 1f, g). Expressed in cells, caGEQO showed similar emission spectra as *in-vitro* and very little bleaching in time course experiments (Extended Data 1c, d).

### GEQO sensors for different analytes and subcellular compartments

The linear design of the GEQO sensor family deviates from the nested construct design of the related MatryoshCaMP^23^ sensors and features a number of restriction sites which enable the straightforward exchange of sensor, reference FP and linker regions, as well as the introduction of organelle targeting motifs (Fig. 1c). By incorporation of different CAAX-motifs^30^, we generated a series of plasma membrane targeted caGEQO calcium sensors (Fig. 1c, Extended Data 1e). The H-Ras derived CAAX-motif resulted in the most consistent plasma membrane localisation enabling the assessment of local changes in calcium concentration in the immediate vicinity of the plasma membrane (Fig. 1c, Extended Data 1e). By exchanging the jGCaMP8f FP with the iATPSnFR^8^ and cAMPing1^6^ biosensors, we generated the ATP and cAMP sensitive atpGEQO and campGEQO sensors (Fig. 1c). Sensitivity to ATP and cAMP was established by monitoring the decrease of ATP concentration induced by pharmacological inhibition of glycolysis and oxidative phosphorylation^31,32^ (Fig. 1c) and the increase of cAMP in response to Forskolin treatment^33^, respectively (Fig. 1c). Taken together, we demonstrate that existing green intensiometric biosensors can be converted by straightforward restriction cloning into ratiometric GEQO sensors for quantitative analyte determination in individual subcellular compartments.

### Absolute quantification and parametrisation of single-cell calcium transients

We next developed a workflow to derive absolute calcium and cAMP concentrations from time resolved GEQO data in single cells. Briefly, we acquired a set of calibration curves on a point scanning confocal microscope (Olympus FluoView3000) using purified recombinant caGEQO protein over a wide range of protein and calcium concentrations, employing the same settings used for imaging cells (Fig. 2a). The fluorescence intensity of the reference channel was related to known caGEQO concentrations in a calibration curve, which enabled us to convert reference fluorescence intensities of individual cells into absolute caGEQO expression levels. By relating the fluorescence intensity of the sensor channel with both known calcium and caGEQO concentrations, we determined the fraction of calcium bound sensor (Θ) for a broad range of calcium concentrations. This yielded a binding curve which was used to determine the concentration of free calcium in solution (Figure 2a, see Supplementary Information for details). Reassuringly, the calcium binding affinity was found to be very similar to the value obtained in bulk plate reader experiments (105 ± 1.4 nM vs 124 ± 0.9 nM) indicating the robustness of the approach (Fig. 2a, Extended Data 1f, g). To probe how external ligand concentrations are translated into intracellular signalling by G-protein coupled receptors (GPCRs), we acquired time trace data of calcium transients induced by varying Neurotensin (Nts) concentrations in HCT116 cells^34^ (Fig. 2b). Fluorescence time trace data were bleach corrected using control time series experiments of untreated cells (Extended Data 1d). Using the parameters determined by the microscope calibration, bleach corrected fluorescence intensities were converted to obtain time trace data expressed in absolute concentrations (Fig. 2c, Extended Data 2, see Supplementary Information for details).

**Figure 2.**
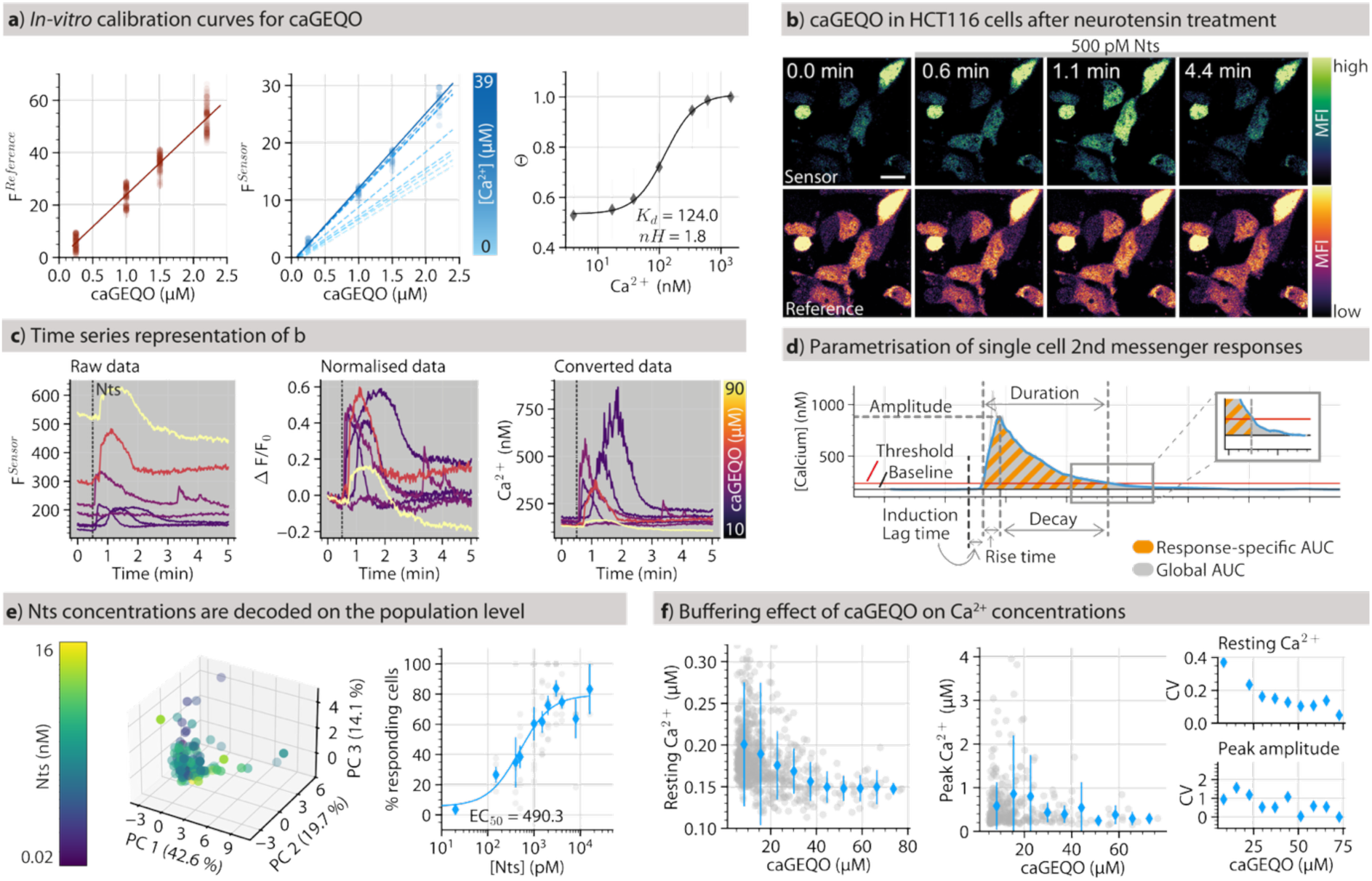
Absolute quantification alleviates analyte buffering. **a)** Three calibration curves (i, ii, iii) are used to convert fluorescence intensities (FI) to absolute Ca^2+^ concentrations. (i) relates the reference channel fluorescence (𝐹^𝑅𝑒𝑓𝑒𝑟𝑒𝑛𝑐𝑒^) to the concentration of purified caGEQO protein. (ii) relates the caGEQO concentration to the sensor channel fluorescence intensity (𝐹^𝑆𝑒𝑛𝑠𝑜𝑟^) at saturating calcium conditions (dark blue, solid line; dots indicate individual measurements. Six lower calcium concentrations shown in light blue, dashed lines). (iii) relates 𝛩 to the calcium concentration (mean ± sem is shown with fitted binding curve). Data from three independent experiments. **b)** Time lapse montage showing HCT116 cells transiently expressing caGEQO treated with 100 nM neurotensin (Nts) after 30 s of imaging (top row: sensor channel in green, bottom row: reference channel in inferno; Scale bar: 25 µm). **c)** Time traces of 6 exemplary cells in (b) showing different representations of the same data: Left: unprocessed sensor fluorescence; middle: normalised sensor fluorescence (ΔF/F_0_); right: absolute Ca^2+^ concentrations over time (min). The expression level of caGEQO is indicated for each time series by a colour value from the inferno colour map. **d)** Parameters extracted from individual single cell calcium concentration profiles as a function of time **e)** Principal component analysis (PCA) of Ca^2+^ responses of single cells (n = 131) to different Nts concentrations (explained variance of each PC indicated in %, each dot represents the Ca^2+^ responses of a single cell; Nts concentration is indicated in colour). Percentage of responding cells at different Nts concentrations (right; grey dots represent independent experiments, blue markers show the mean ± 95% confidence interval (bootstrapped), blue line represents a dose-response fit; n = 614). **f)** The resting concentration of Ca^2+^ (left) and the peak amplitude of Ca^2+^ responses (middle) over different expression levels of caGEQO (Grey dots represent individual cells, blue markers show mean and standard deviation of GEQO concentration bins) and their covariance (CV)

We then defined a parameter set designed to capture the unique properties of single cell time trace data expressed both in absolute concentrations and normalised formats (relative quantification, ΔF/F_0_, ΔR/R_0_) to comprehensively assess differences in second messenger responses on a single cell level (Fig. 2c). Specifically, we quantified features describing each cells’ calcium response by its duration, peak amplitude, rise time, decay time, area under the curve (globally and specifically during responses), number of responses, resting calcium level/concentration, and elapsed time to response onset (Fig. 2d).

Principal Component Analysis (PCA) helps simplify complex, high-dimensional data by projecting it onto two- or three-dimensional plots, where each dot represents the calcium response of a single cell. Beyond visualization, PCA also reveals how the individual quantified features influence the data distribution. This is done using vectors that indicate both the direction and strength of each feature’s contribution in principal component space. To further analyse calcium responses, we apply K-means clustering, a method that groups similar responses together. We use the silhouette score as a metric to assess clustering quality and determine the number of clusters^35^. The score is computed as an average for all clusters and ranges from −1 to 1, with higher values indicating better-defined clusters. This approach allows us to compare calcium responses both at the single-cell and population level.

Using these methods, we investigated whether HCT116 cells can distinguish between different concentrations of neurotensin (Nts), the natural ligand of the NTSR1 receptor, which these cells express endogenously. We found that calcium response dynamics and amplitudes on the single cell level were not correlated with Nts stimulus strength despite widely varying parameters for individual time traces (Extended Data 3). In contrast, the population level analysis showed a classical dose-response curve with an EC_50_ of 490 pM (literature values 150-1800 pm^36,37^; Fig. 2e). This implies that information processing during neurotensin-induced calcium signalling likely does not rely on transient dynamics but follows a binary logic, at least in HCT116 cells. Taken together, our approach allows to capture single cell calcium responses in a compact set of parameters and identify characteristic patterns in response dynamics or the absence thereof.

### GEQO sensors alleviate artifacts caused by analyte buffering

By using genetically encoded fluorescent biosensors researchers always assume the risk of perturbing the investigated process due to analyte buffering. For second messengers, this can result in lower baseline concentrations, dampened peak amplitudes, and overall altered signalling dynamics. These artifacts are widely known and have been extensively discussed in the literature^38–42^, but so far it has been difficult to quantify and avoid them. We reasoned that our GEQO platform offers a straightforward way to directly assess analyte buffering and identify suitable sensor expression levels for live cell applications.

We first compared baseline cytosolic calcium concentrations in HCT116 cells transiently expressing caGEQO under a CMV promoter; a widely used approach for introducing biosensors into target cells^43^ (Fig. 2f). We found that increasing sensor concentrations were associated with a decrease in baseline calcium concentrations (192 ± 66 nM for cells expressing 5-12 µM caGEQO to 138 ± 7 nM for cells expressing 76-85 µM caGEQO; Fig. 2f, left panel). A similar dampening effect was found for the amplitude of Nts induced calcium transients. In particular, high amplitude transients all but disappeared at higher caGEQO expression levels (Fig. 2f, middle panel). Both effects are highly likely a direct consequence of calcium buffering, as is indicated by decreasing coefficients of variation with increasing sensor concentration, which is a hallmark of buffering^44^ (Fig. 2f, right panel). Based on these data, we propose that the intracellular concentration of GCaMP type calcium sensors should be kept below 20 µM and ideally lower.

### Sensor concentration differentially affects PKA-based cAMP biosensor responses

Using campGEQO, we next assessed the effect of sensor concentration on fluorescent readout in cAMP measurements. campGEQO uses the green intensiometric cAMPing1 sensor as the analyte-sensitive protein domain, which is derived from the protein kinase A regulatory subunit 1α (PKA-R1α), that features two cAMP binding sites. The fluorescent protein is inserted into the cyclic nucleotide binding site A (CNB-A), the higher affinity second binding site CNB-B is left intact.^45^ This means that the expected fluorescence response to changes in cAMP concentration does not follow a simple binding model of two interacting species, which complicates absolute quantification. With increasing sensor concentrations, the expected midpoint of the fluorescence response is more sensitive to increasing sensor concentration compared to a sensor design that uses only one ligand binding site (Fig. 3a, see Supplementary Information for detailed discussion). Using purified full length cAMPing1 protein, as well as a truncated version that lacks the CNB-B site (cAMPing-ΔB) we confirmed that the CNB-B site indeed affects the sensor response as predicted (Fig. 3b). The ratiometric campGEQO behaved like a single-binding site sensor with regard to the concentration dependent fluorescence response midpoint shift, offering overall preferable sensor properties despite a slightly reduced dynamic range.

**Figure 3.**
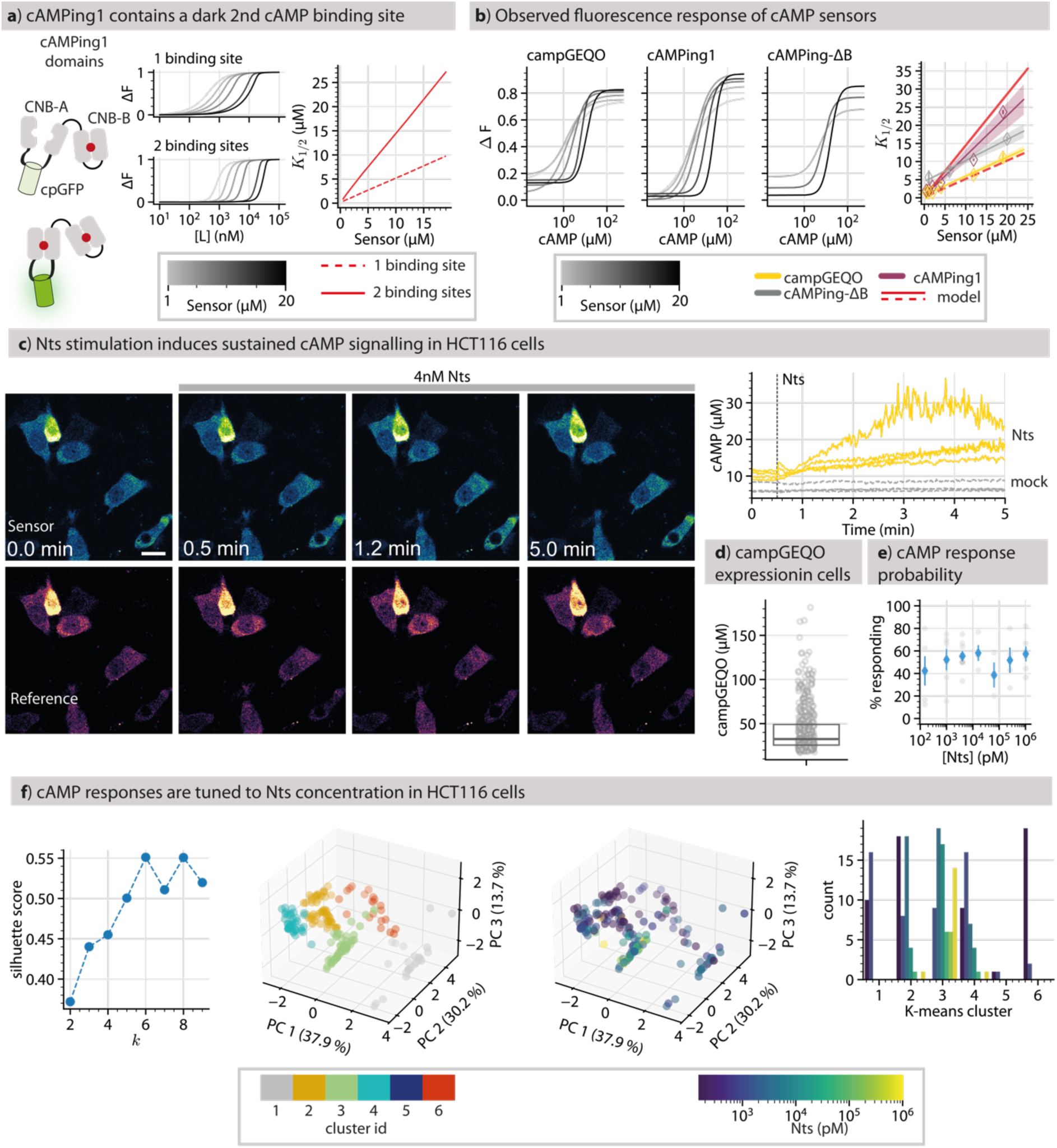
campGEQO reveals cAMP responses tuned ligand concentration. **a**) cAMPing1 has two binding sites for cAMP, only the conformation of CNB-A has an effect on fluorescence (left panel). When the sensor concentration is increased, the fluorescence response shifts to higher ligand concentrations (middle panel, sensor concentration in greyscale), which is illustrated as a shift in the midpoint (K_1/2_) of the fluorescence response (right panel; regression line of midpoint shift for one binding site is dashed, for two binding sites solid). **b**) Fluorescence response of campGEQO (left), cAMPing1 (middle), and cAMPing-ΔB (right) at different sensor concentrations (greyscale). Regression lines (right) show the observed change in K_1/2_ for each construct (cAMPing1: purple, cAMPing-ΔB: grey, campGEQO: yellow, markers show mean K_1/2_ ± sem) The predicted change in K_1/2_ for one (dashed) and two (solid) binding sites is shown in red. **c**) Time lapse of HCT116 cells treated with 4 nM neurotensin (Nts; top: sensor channel in green, bottom: reference channel in inferno LUTs) and exemplary time traces (right panel) of Nts (4 nM, yellow) and mock (grey) treated cells. **d**) Quantified expression level of campGEQO in HCT116 cells (dots represent individual cells, box extends from the 25th to the 75th percentile, horizontal line indicates the median). **e**) Percentage of responding cells at different Nts concentrations (grey dots represent independent experiments, blue markers show the mean ± 95% confidence interval (bootstrapped, n = 329), **f**) cAMP responses (n = 142) of HCT116 cells treated with different concentrations of Nts were grouped into 6 clusters, based on silhouette score (first panel). The distribution of cells in principal component space is shown with cluster id (second panel, discrete colours) and Nts concentration (third panel, viridis colourmap) indicated. Fourth panel shows distribution of Nts concentration used by cluster id.

We used the knowledge of intracellular sensor concentration to analyse cAMP responses on a single cell level. We stimulated cAMP responses in HCT116 cells using a range of Nts concentrations and analysed only cells expressing campGEQO between 15 and 30 µM to eliminate biases from shifting sensor concentrations. In contrast to Nts induced calcium signalling, cAMP signalling did not follow a dose response profile on the population level, with regard to the percentage of responding cells (Fig. 3e). Instead, single cell responses to low and high Nts concentrations clustered in different populations. While cells treated with low Nts concentrations where distributed over multiple clusters, treatment with higher Nts concentrations resulted in more similar cAMP responses, mostly grouped to cluster 3 (Fig. 3f). These data indicate that the same receptor (NTSR1) can transduce information either as a binary on/off signal as seen in calcium responses, or encoded in the single cell dynamics of cAMP responses.

Taken together, these data demonstrate that artifacts due to analyte buffering can be minimized by using GEQO sensors. Furthermore, certain flaws in original sensor design are readily observed and can be accounted for by adapting the analysis pipeline.

### Absolute quantification of calcium dynamics prevents information loss

Calcium signalling of varying temporal dynamics is a key feature of organismal development. Cells encode information about receptor-ligand concentrations, metabolic state, and tissue context in amplitude, frequency, and shape of calcium transients. Processing of this information ultimately influences cell fate decisions^46–48^. We thus asked whether quantitative calcium imaging using GEQO sensors can capture information encoded in calcium response patterns, which might be lost during relative quantification of intensiometric and ratiometric datasets. GEQO sensors can be used as intensiometric (sensor channel only) or traditional ratiometric (sensor/reference channel ratio) biosensors. Therefore intensiometric, ratiometric, and absolute quantified time traces can be obtained from the same GEQO data set and used as input for a comparative parametrisation analysis.

In a first application, we used human pancreatic progenitor (ePE) cells^49^ derived from the embryonic stem cell line H1 and transfected with caGEQO. Calcium responses were induced using either ATP or arachidonic acid, based on known receptor expression patterns^50,51^. The percentage of responding cells did not differ much on the population level and the amplitudes of the average traces was also similar (Fig. 4a). Analysis of single cell traces showed that ATP and arachidonic acid treated cells form two distinct populations in PCA, which are primarily separated along the PC2 axis, irrespective of data type (Fig. 4b-c). However, the contribution of the underlying features differed strongly between intensiometric, ratiometric, and absolute quantification data sets. Generally speaking, the parameters can be grouped into three classes: features that capture response shape (rise and decay times, response width), features that capture response initiation and frequency (lag time and peak number) and features that capture response intensity (peak amplitude and released calcium).

**Figure 4.**
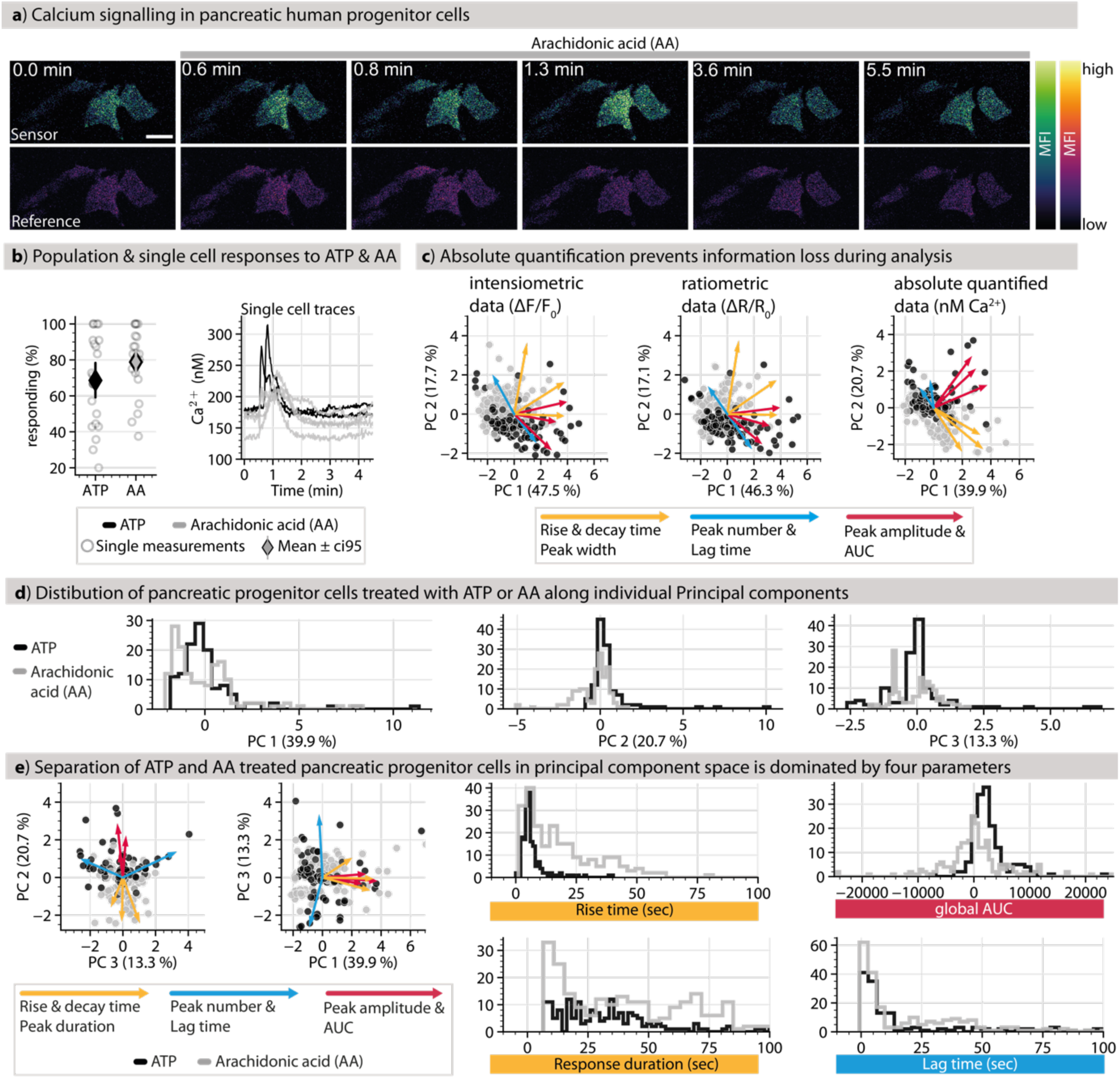
Absolute quantification of Ca^2+^ signalling enhances conclusiveness of data. **a)** Time lapse montage of human pancreatic progenitor (ePE) cells treated with arachidonic acid (AA) after 30 s (top: sensor channel in green, bottom: reference channel in inferno; Scale bar: 25 µm). **b)** Probability of Ca^2+^ response of ePE cells to AA (n = 282) and ATP (n = 223; left panel, circles represent independent experiments, diamond markers represent mean ± 95% confidence interval) and 4 exemplary responses of cells treated with either AA or ATP (right panel). **c)** PCA of single cell Ca^2+^ response profiles (n = 505) evaluating the same data set as intensiometric (left panel) and ratiometric (middle panel) data or as absolute Ca^2+^ concentrations (right panel; dots represent individual cells, loading vectors shown in colour; explained variance indicated in percent). **d)** Histograms showing the distribution of pancreatic progenitor cells treated with arachidonic acid (AA, grey) or ATP (black) along individual principal components. **e)** Distribution of single cell Ca^2+^ response profiles to AA (grey) and ATP (black) treatment along the 2^nd^ & 3^rd^ (left panel) and along the 1^st^ & 3^rd^ principal component (PC, right panel). Loading vectors are shown colour coded for three groups of parameters: i: rise & decay times, response duration (yellow); ii: peak number & lag time (blue); iii: peak amplitude and response specific & global area under the curve (AUC, red). Histograms show the distributions of four exemplary parameters, rise time, response duration, global AUC, and lag time. The explained variance of each PC is indicated in percent.

For intensiometric and ratiometric data, feature contribution to response distribution in principal component space did not exhibit a readily apparent pattern (Fig. 4c left & middle panel). In the quantified data set, parameter groups describing response shape, initiation and frequency or response intensity contribute in distinctly different ways to the distribution of individual responses in PC space (Fig. 4c right panel). Overall, features capturing response shape and intensity features dominated the separation of AA and ATP induced responses along the first and second principal component axis (Fig. 4c-e).

In a second case study, we used caGEQO to study spontaneous calcium transients in developing zebrafish (*Danio rerio*) embryos, specifically monitoring transients known to occur in the skin epithelium (Fig. 5a, b). We therefore generated a stable line expressing caGEQO under a *ubiquitin* promoter with a cytoplasmic biosensor concentration of ∼351 ± 1.5 nM (Fig. 5b), ensuring that sensor buffering should be limited. Calcium transients were captured by time lapse imaging and analysed as intensiometric, ratiometric, and quantified data sets (Fig. 5c). We found that the contribution of parameters to the distribution of responses in PC space mirrored the trends observed during the analysis of the human pancreatic progenitor cell datasets. Specifically, the separation along the PC1 and PC2 axes was primarily due to features capturing response shape (rise and decay times and response duration) and intensity (amplitude) in the quantified dataset, whereas no such pattern was found for the intensiometric and ratiometric datasets (Fig. 5c). The two investigated systems are from evolutionary distant organisms and the acquired traces were rather different (mostly single peaks for pancreatic progenitor cells vs. oscillatory patterns in the zebrafish epithelium). In both cases, information was captured in quantified datasets that was lost in both intensiometric and ratiometric datasets implying that absolute quantification of single cell calcium transients using GEQO sensors offers significant benefits compared to established data acquisition routines using other types of biosensors.

**Figure 5.**
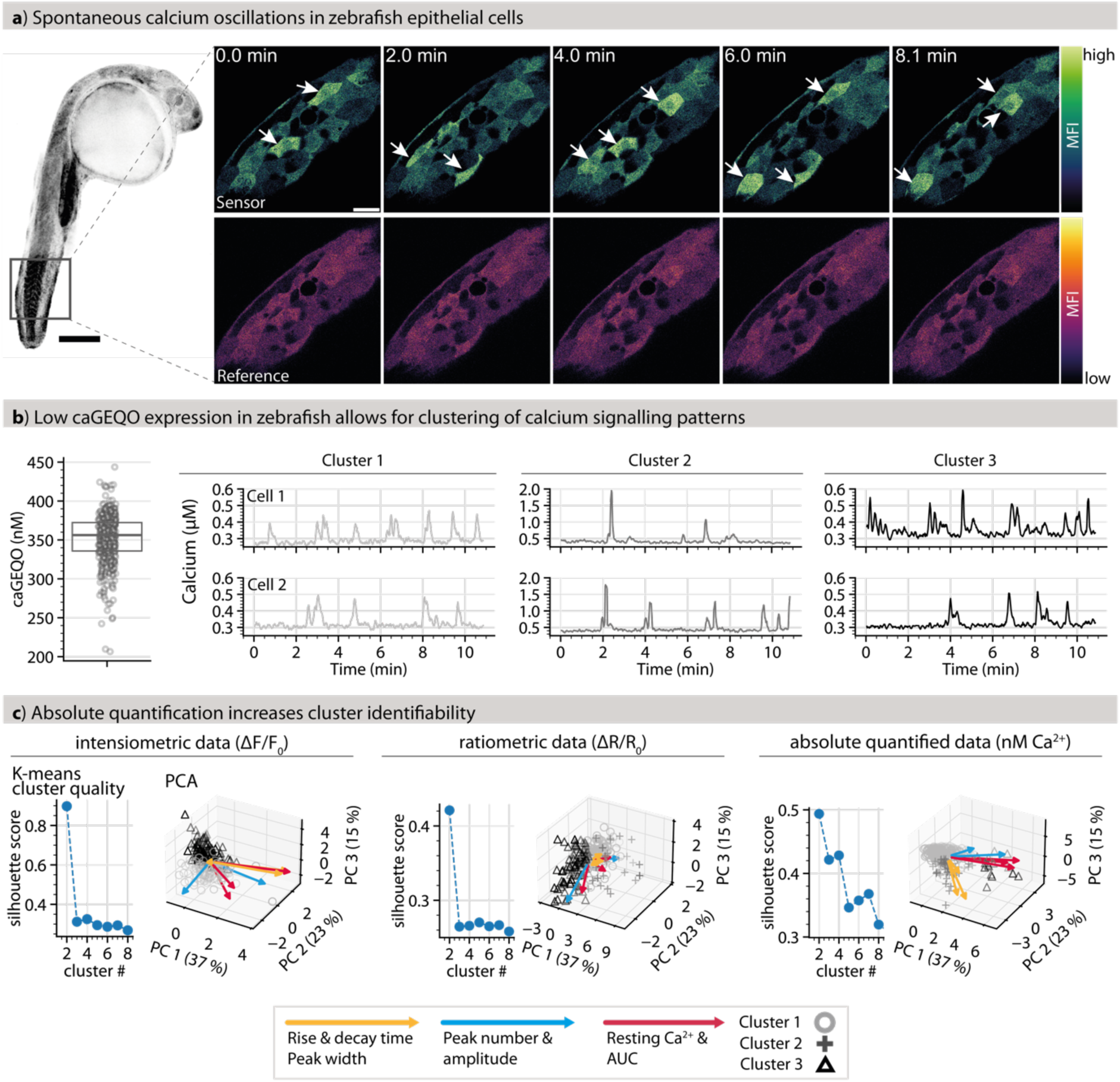
Absolute quantification of Ca^2+^ signalling in zebrafish embryos reveals patterned data. **a)** *(left)* Transgenic zebrafish embryo at 24 hours post fertilisation (hpf), expressing caGEQO. Scale bar: 250 µm. Black inset highlights region of interest imaged at higher resolution to the right. *(right)* Time lapse images of the apical epithelial cells of caGEQO transgenics. caGECO sensor channel in green hot and reference channel in inferno LUTs. Scale bar: 25 µm. **b)** caGEQO concentration in zebrafish epithelial cells (n = 432) driven by expression of the *ubiquitin* promoter, at 24 hpf (*left*). Points represent individual cells, boxes extend from the 25th to the 75th percentile, horizontal line indicates the median. Representative calcium concentration along time of three clusters identified by K-means clustering. Open circles, light grey: cluster 1; crosses, dark grey: cluster 2; open triangles, black: cluster 3. **c)** Evaluation of the K-means clustering by silhouette score for 2-9 clusters and PCA of single cell Ca^2+^ response profiles from acquired zebrafish time lapses, evaluating the same dataset as intensiometric (left), ratiometric (middle), and absolute Ca^2+^ concentrations (right; points represent individual cells formatted by cluster identity. Open circles, light grey: cluster 1; crosses, dark grey: cluster 2; open triangles, black: cluster 3; loading vectors are shown in colour).

To fully explore the information content of absolute quantified datasets, we sorted the traces within all zebrafish datasets into three clusters using K-means clustering^52^. In case of the intensiometric dataset, only two clusters were meaningfully populated, whereas the third contained a single trace. This is also reflected by the silhouette score, a measure of clustering quality^35^, which suggests that the intensiometric traces can be meaningfully sorted into maximum two clusters (Fig 5c, left panel). While for ratiometric data all three clusters are populated, the silhouette score rapidly decreases from two to three clusters in this case as well, suggesting no meaningful separation (Fig 5c, middle panel). For absolute calcium concentrations, the silhouette score remains low but transiently increases for three and four clusters. This pattern suggests competing structures or hierarchical relationships within the data, which may be lost when assuming fewer clusters. In principal component space, the three clusters align with the directions of the loading vectors. This alignment suggests that the clustering reflects intrinsic data structure, with the loading vectors appearing to group into three distinct classes (Fig 5c, right panel).

For both the zebrafish and pancreas progenitor cell data sets, the response features derived from calcium time trace data contribute differently to principal component analysis for relative and absolute quantification approaches. Taken together, this analysis of calcium dynamics in different physiologically relevant systems demonstrates that absolute quantification of second messenger transients is necessary for a faithful assignment of the causal features in statistical data analysis and clustering approaches.

## Discussion

We here report the GEQO platform of quantitative genetically encoded fluorescent biosensors, which enables straightforward, direct quantification of analyte concentrations by fluorescence microscopy in live cell experiments. The sensor design allows for converting existing intensiometric GFP-based biosensors to into quantitative GEQO sensors. By monitoring calcium and cAMP levels in cell and tissue assays, we demonstrate that (i) GEQO sensors alleviate artifacts commonly associated with fluorescent biosensors, such as analyte buffering; and (ii) that statistical analysis of quantified calcium responses captures information which is lost in intensiometric and ratiometric data sets.

Compared to existing biosensor-based approaches for quantifying analytes in living cells, GEQO sensors offer a number of advantages. GEQO sensors retain most of the dynamic range and signal-to-noise ratio of intensiometric sensors, which constitutes an advantage over FRET sensors, in particular for single cell measurements. FLIM sensors allow for the direct determination of analyte concentrations through fluorescence lifetime measurements, whereas GEQO sensors require instrument calibration. Advantages of GEQO sensors are the wide applicability, as they do not require specialised microscopes, and the straightforward quantification of both intracellular sensor and analyte concentration. The quantification of sensor expression levels allows for estimating analyte buffering by the sensor and provides a pathway to correct for artifacts caused by varying sensor levels. Taken together, GEQO sensors enable imaging based workflows across many platforms that are equivalent to widely used mass spectrometry approaches, e.g. for quantitative biomarker detection^53^.

Absolute quantification of cellular signalling dynamics by straightforward fluorescence microscopy will have major benefits for cell and developmental biology, in particular taking into account that the GEQO platform can be readily expanded to include additional analytes. Furthermore, the possibility of standardised workflows is the prerequisite for applications in diagnostics^54^, for instance after delivery of biosensors via lipid nanoparticles into patient derived organoids^55,56^. We expect that GEQO sensors will be widely used due to their adaptability, compatibility with a broad range of imaging setups, and ease of application.

## Acknowledgements

AN and SMK gratefully acknowledge financial support by the European Research Council (ERC) under the European Union’s Horizon 2020 research and innovation program (grant agreements no GA 758334 ASYMMEM). E.N., S.K., and R.M. acknowledge funding from the Max Planck Society and the Deutsche Forschungsgemeinschaft (DFG, German Research Foundation) under Germany’s Excellence 381 Strategy -EXC-2068-390729961 - Cluster of Excellence Physics of Life of TU Dresden. JL acknowledges funding by the DFG under the Walter Benjamin programme (grant number: 523926206). AGB acknowledges funding by the Max Planck Society. We thank the following services and facilities at MPI-CBG Dresden for their support: the Light Microscopy Facility, the Protein Biochemistry Facility, and the Technology Development Studio. We thank Yung Hae Kim for significant strategic support for the transfection of human ePE cells. We thank Jana F. Fuhrmann for outstanding support and expert advice on data analysis and critical feedback on the manuscript and Michele Marrass for his expert advice. We particularly thank Joachim Goedhart for his important feedback on the manuscript.

## Author contributions

SMK, EN, and PB prepared samples and acquired the datasets. JFL and MBA prepared samples. SK generated *D. rerio* lines. SMK and EN analysed imaging data. SMK developed the quantification and parametrisation workflow and analysed the data. SMK and AN designed the project. AN, EG, RM, and AGB supervised research. SMK and AN wrote the manuscript. All authors read and commented on the manuscript.

## Data Availability

The raw data and code to analyse all data can be accessed at: https://doi.org/10.17617/3.9IHKTD

---

**Extended Data Figure 1.**
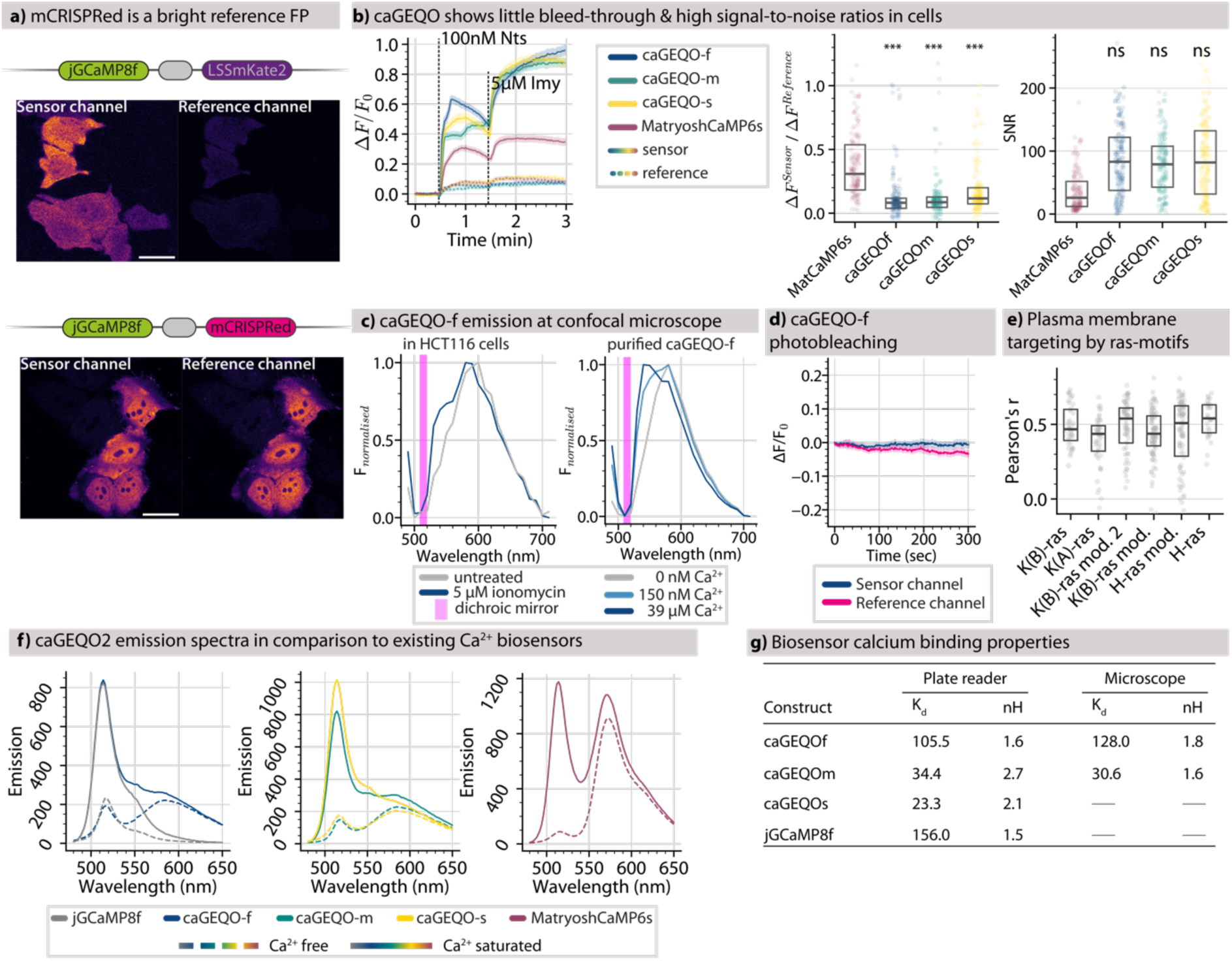
Development of GEQO sensors. **a)** Comparison between LSS-mKate2 based GEQO1 (top) and mCRISPRed based GEQO2 (bottom). The linear construct design is shown and representative images of the sensor and reference channels from HCT116 cells transiently expressing either construct (Scale bar: 50 µm). **b)** Average time traces of HCT116 cells transiently expressing caGEQO-f (blue, n = 154), -m (green, n = 141), -s (yellow, n = 171), or MatryoshCaMP6s (purple, n = 132) showing the sensor channel as solid and the reference channel as dashed lines (left; shaded areas indicate the standard deviation of the mean). Cells are treated with 100 nM neurotensin (Nts) after 30 s and 5 µM ionomycin (Imy) after 90 s. The fluorescent bleed-through and signal-to-noise ratio quantified from live-cell experiments are shown in box-stripplots (right; Boxes extend from the 25th to the 75th percentile, horizontal lines indicate the median, each measurement is represented by a dot. P-values calculated using the Welch t-test). **c)** Left panel: Emission spectra of caGEQO-f in HCT116 cells before (grey, n = 67) and 5 minutes after ionomycin treatment (blue, n = 104). Right panel: emission spectra of purified caGEQO-2 in the presence of 0 nM (grey), 150 nM (lght blue), and 39 µM (dark blue) free calcium. Magenta line indicates the non-transmissive spectrum of the dichroic mirror used for imaging. **d)** Photobleaching of the caGEQO-f in untreated HCT116 cells; sensor channel (blue) and reference channel (magenta), n = 115. **e)** Quantification of the membrane targeting efficiency by CAAX motifs derived from different ras proteins as the Pearson’s correlation coefficient (Boxes extend from the 25th to the 75th percentile, horizontal lines indicate the median, each measurement is represented by a dot, n ≥ 32). **f)** Emission spectra of caGEQO2f (blue) overlaid with jGCaMP8f (grey, left), caGEQO2m (green) overlaid with caGEQO2s (yellow, middle), and MatryoshCaMP6s (purple, right). Calcium free spectra are represented by dashed lines, calcium saturated spectra (39 µM) by solid lines. **g)** Table summarising the *in-vitro* calcium binding properties of caGEQO-f, -m, -s, and jGCaMP8f measured using a plate reader or during microscope calibrations.

**Extended Data Figure 2.**
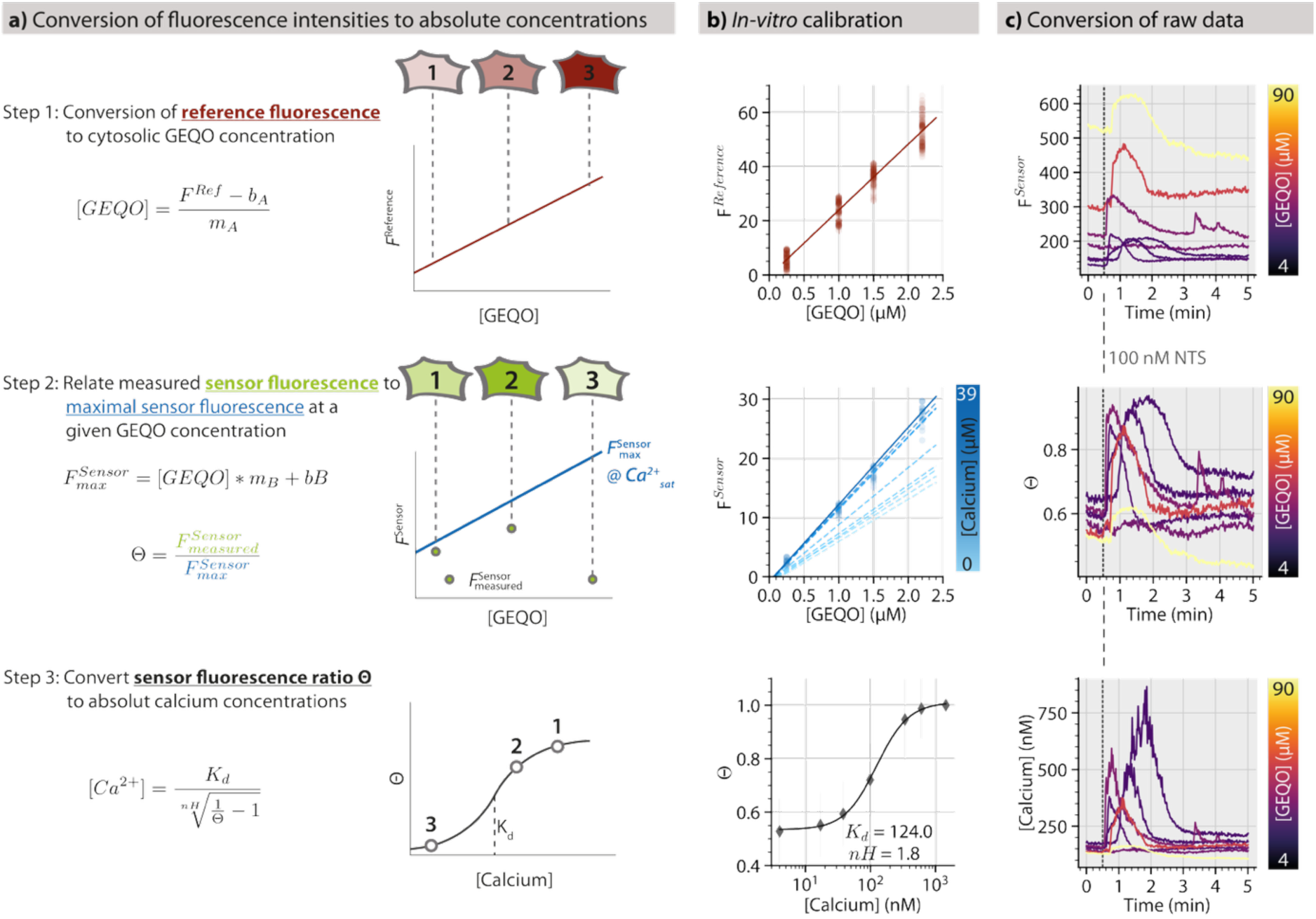
Conversion of fluorescent intensities to absolute concentrations. **a)** The schematic workflow used to convert fluorescence intensities to absolute concentrations of calcium and GEQO concentration. Cells with different expression levels of GEQO protein, as indicated in shades of red representing the fluorescence in the reference channel. Calibration curve A (red line), relating reference fluorescence to sensor concentration, is used to determine the GEQO concentration. The same three cells showing different fluorescence intensities in the sensor channel as represented by different shades of green. Calibration curve B (blue line) relates the sensor channel fluorescence to sensor concentration at saturating calcium concentrations and is used to estimate the maximum sensor channel fluorescence for any given GEQO concentration. The ratio of the measured sensor fluorescence and the expected maximum sensor fluorescence at a given GEQO concentration yields Θ. Calibration curve C (grey line) relates Θ to the concentration of free calcium. Using the parameters K_d_ (the calcium concentration producing half maximum Θ values), nH (the Hill coefficient), and the Θ values estimated using calibration curve B, the free calcium concentration for each cell can be estimated. **b)** Calibration curves A, B, and C as measured using purified caGEQO2f. **c)** Time traces of representative HCT116 cells - transiently expressing caGEQO2f and treated with 100 nM neurotensin (NTS) after 30 seconds – at different stages of conversion (GEQO expression level indicated in colour). Top: fluorescence intensity of the sensor channel. Middle: The same time traces normalised to Θ values using calibration curve B. Bottom: The same time traces converted to absolute calcium concentrations using calibration curve C.

**Extended Data Figure 3.**
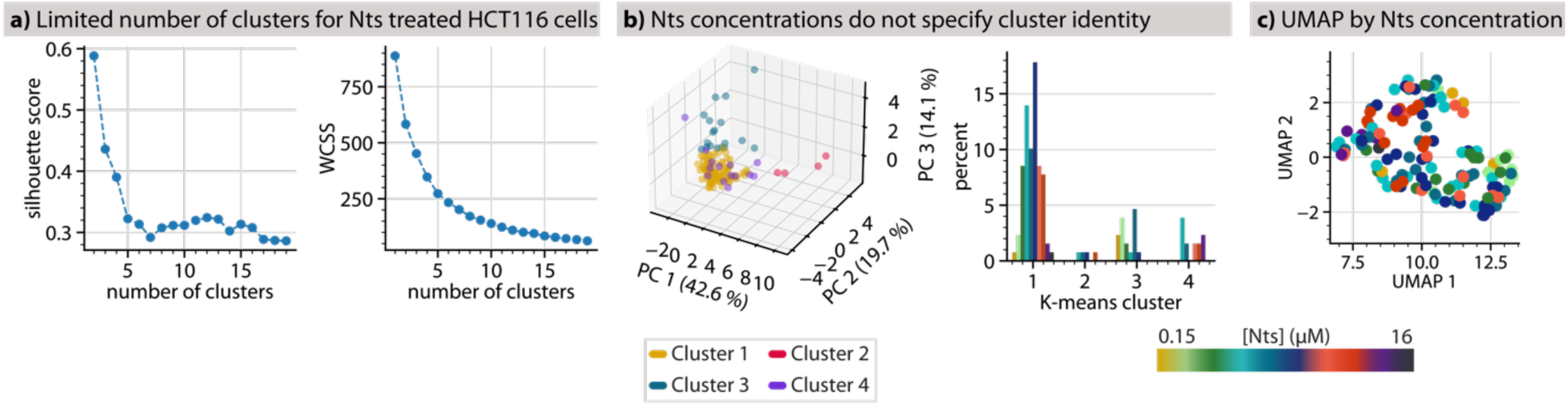
Clustering of neurotensin mediated calcium responses. **a)** Silhouette score and within cluster sum of squares (WCSS) at increasing numbers of K-means clusters for neurotensin (Nts) treated HCT116 cells (n = 771) transiently expressing caGEQO (Fig. 2) **b)** PCA of single cell Ca^2+^ response profiles to Nts, colour coded by K-means cluster identity (left). Percentage of cells responding to the indicated Nts concentrations within each K-means cluster (right). **c)** Distribution of Ca^2+^ response profiles after dimensionality reduction using Uniform Manifold Approximation and Projection (colour coded by Nts concentration).

