## Supplementary information for "GEQO biosensors for absolute analyte quantification in single cells"

**Material & Methods**

**Ethics statement**

This study followed European Union animal welfare directives (2010/63/EU) and German law applied to zebrafish work, with license #TVV52/2021 - ‘Generierung von Zebrafischlinien zur Untersuchung der Größe und Form von Organen und Organellen‘, project leader Rita Mateus. Genetic engineering work was carried out in an S1 area (MPI-CBG, S1-Labore 4., Az.: 54-8451/103, project leader Rita Mateus), following guidelines according to Section 21, Paragraph 1 of the German Genetic Engineering Act, and within projects 01 and 03 from the Mateus laboratory, according to Section 28 of the German Genetic Engineering Safety Ordinance (GenTSV).

**DNA constructs**

GEQO1f constructs were designed *in-silico* to include a multiple cloning site of 6 unique restriction sites (NcoI, NotI, EcoRI, BamHI, AscI, and HindIII) with jGCaMP8f flanked by the NotI & EcoRI sites, a flexible linker flanked by EcoRI & BamHI, and LSS-mKate2 flanked by BamHI & AscI. This construct was procured as a clonal gene in a mammalian expression vector from Twist Bioscience (pTwist-CMV). jGCaMP8m, jGCaMP8s, mCRISPRed, cAMPing1, and iATPSnFR were procured from Twist bioscience as DNA fragments flanked by the correct restriction sites. These fragments were amplified using Twist Primer Set 2. mCRISPRed was used to replace LSS-mKate2 in the BamHI-AscI site to generate GEQO2f. jGCaMP8m and s were used to replace jGCaMP8f to generate GEQO2m and GEQO2s, respectively. campGEQO and atpGEQO were generated by inserting mCRISPRed into the NotI-EcoRI site, then inserting cAMPing1 and iATPSnFR into the BamHI-AscI site, respectively. For expression in insect cells, the finished constructs were transferred from pTwist-CMV to pOCC5[1] using NotI and HindIII. Insect cell expression constructs of MatryoshCaMP6s, jGCaMP8f, cAMPing1, cAMPing1-ΔB, and CNB-B were produced by procuring fragments flanked with the according restriction sites from Twist Bioscience and inserting them into pOCC5. For zebrafish constructs, all GEQO2 variants were PCR amplified from the above constructs using primers containing attB1 and attB2 recombination sites (fwd: 5'-GGGGACAAGTTTGTACAAAAAAGCAGGCTTAatgggaacgcgtcgca-3', rev: 5'-GGGGACCACTTTGTACAAGAAAGCTGGGTTttacggcgcgccct-3'). GEQO2 fragments were recombined with pDONOR221 (#208 from the Tol2 Kit) using Gateway BP-cloning (BP clonase II Enzyme-Mix Invitrogen #11789020) to generate Middle Entry (pME) clones. The obtained pME clones were each then recombined in LR reactions (LR Clonase II Plus Enzyme from Invitrogen #12538120), using pENTR5'_ubi (addgene #27320), containing the ubiquitin (ubi) promoter, and pDEST-pTol2pA R4-R2 backbone (#465 from the Tol2 kit), to obtain the T2-ubi:GEQO2-1f, T2-ubi:GEQO2-1m and T2-ubi:GEQO2-1s plasmids.

**Protein expression and purification**

Proteins were purified from Sf9 or Tni insect cells. For protein expression fresh batches baculovirus were produced for each construct according to the FlexiBac protocol[1]. In brief: Sf9 insect cells were co-transfected using linearised viral genome and expression plasmid of the respective construct. P1 virus from transfected cells was used to infect fresh Sf9 cells to produce P2 virus. P2 virus was used to infect 500 ml of Sf9 or Tni cells ($10^6$ cells/ml in ESF921 medium) and cells were harvested by centrifugation after three days. Cell pellets were resuspended in 50 ml buffer (2x PBS, 1 mM DTT (Thermo Scientific™ R0861), 20 mM Imidazole (Sigma Aldrich 56750), 0.25U/ml Benzonase, 1 mM $MgCl_2$, 1x protease inhibitor cocktail (Roche 05056489001)), mechanically lysed using an LM20 Microfluidizer® (at 20 000 PSI, Microfluidics International Corporation), and centrifuged for 1 h at 4100 rpm and 4 °C to remove cellular debris. The lysate was loaded onto 5 ml HisTrap® immobilised metal affinity chromatography (IMAC (Cytiva 17524801) columns, washed with 10 column volumes (CV) low salt buffer (2x PBS, 20 mM Imidazole, 1 mM DTT), 10 CV high salt buffer (6x PBS, 1 mM DTT), and 10 CV low salt buffer. Column bound protein was eluted in 2 ml 250 mM Imidazole in 2x PBS, 1 mM DTT. Protein was then dialysed overnight at 4°C against 2x PBS, 1 mM DTT while cleaving the 6xHis-tag using 6xHis-tagged 3C protease.

Remaining impurities and the 3C protease were removed the following day using a 5 ml HisTrap® column, collecting the flow through. Purified protein concentration was adjusted to ~500 μM according to NanoDrop measurements, aliquoted, and snap frozen in liquid nitrogen. Purified protein was quantified using the Thermo Fisher Scientific Pierce™ BCA kit (23225).

**Fluorimetric characterisation of fluorescent sensors**

*In-vitro* fluorimetric characterisation was performed using a fluorescence plate reader (Spark 20M, Tecan). Calcium titrations were prepared by mixing stock solutions of the commercial Calcium Calibration Kit #1 (Invitrogen Life Technology C3008MP) according to the manufacturer's instructions, yielding 11 buffers of different free calcium concentrations. cAMP titrations were prepared by diluting a stock solution of 5 mM cAMP (Sigma-Aldrich A6885) to the desired target concentrations.

10 μl of purified protein and 90 μl of calcium buffer were combined and distributed in triplicates of 30 μl each on a black, flat-bottom 384-well plate (Greiner Bio-One 781900) using a pipetting robot (Freedom Evo20, Tecan). End-point fluorescence emission at $\lambda_{em}$ = 525 nm was recorded using top read mode, 50 nm bandwidth and a gain of 150 at an excitation wavelength of $\lambda_{ex}$ = 445. Binding characteristics were determined by fitting the Hill equation to the measured data. End-point fluorescence spectra were recorded using top read mode, 5 nm bandwidth and a gain of 150. Emission spectra were recorded at $\lambda_{ex}$ = 445. Spectra were background corrected using a buffer control and values of emission maxima were extracted (scipy.measure.find_peaks) to calculate the fluorescent bleed-through between sensor and reference domains as the change in reference channel fluorescence.

**Fluorescence response prediction**

To predict how the fluorescence response is influenced by the concentration of the sensor itself and how it is influenced by the number of binding sites, we estimated the fluorescence response as the concentration of sensor R bound to ligand L as:

$$[LR] = \frac{[L]_{\text{free}} \cdot [R]}{K_d + [L]_{\text{free}}}, \tag{1}$$

Where $K_d$ is the dissociation constant of R. If as second binding site $R_2$ for L is present that binds L with dissociation constant $K_{d2}$, [LR] is dependent on the availability of L which can be calculated as:

$$[L]_{\text{free}} = [L]_{\text{total}} - \frac{[L]_{\text{free}}[R]}{K_d + [L]_{\text{free}}} - \frac{[L]_{\text{free}}[R_2]}{K_{d2} + [L]_{\text{free}}} \tag{2}$$

To predict the fluorescence response LR at a given concentration of R and $R_2$, equation (2) is solved numerically using scipy's root_scalar function. The $[L]_{free}$ is then substituted into equation (1) to solve for [LR] and $[LR_2]$, respectively.

To simulate a sensor with a single binding site, $[R_2] = 0$. For a sensor with two binding sites $[R_2] = [R]$. To simulate the interplay of CNB-A and CNB-B, we assumed the dissociation constants $K_{d\ CNB\_A} = 150$ nM and $K_{d\ CNB\text{-}B} = 4$ nM, as reported by Lorenz et al.[2]

**Mammalian cell culture and imaging**

HCT116 human colorectal cancer cells were grown in McCoy's modified 5A GlutaMax growth medium (Gibco 36600021) supplemented with 10% fetal calf serum (Gibco 10270-106). Transfection of HCT116 cells was performed using X-tremeGene™ HP transfection reagent (Merck XTGHP-RO). Cells were seeded into ibidi 18-well glass bottom µ-slides (81817) at a concentration of 20 000 cells/well, 24 hours before transfection. 600 ng of plasmid DNA were mixed with 1.8 µl of transfection reagent in 200 µl OptiMem (Gibco 31985062) and incubated at room temperature for 15 minutes before adding 800 ml of growth medium and cells were treated with 60 µl/well of transfection mix for 5 hours. Cells were used for experiments after a minimum of 12 hours.

Human expanding pancreatic epithelium (ePE) cells were cultured on fibronectin (Sigma-Aldrich 0895) coated plates in DMEM/F12 medium with Glutamax (ThermoFisher 31331028), supplemented with 64 ng/mL FGF2 (Peprotech 100-18B), 1x B27 (ThermoFisher 17504044), and 10 µM SB431542 (Calbiochem 616464). 10 µM of Rock inhibitor Y-27632 (VWR 688000) were present during the first 24 hours after seeding. ePE cells were used after passage 5. For transfection, ePE cells were seeded into ibidi 18-well glass bottom µ-slides at a concentration of 34000 cells/well, 24 hours before transfection. 100 ng of plasmid DNA were transfected into each well using 0.2 µL of lipofectamine stem transfection reagent (ThermoFisher STEM00001). Cells were used for experiments after 48 hours.

Cells were imaged on an Olympus FluoView3000 point scanning confocal microscope using a 60x (HCT116) or 40x (ePE) oil immersion objective at 37°C, 5% $CO_2$ in imaging buffer (IB: 20 mM HEPES, 115 mM NaCl, 1.2 mM CaCl2, 1.2 mM MgCl2, 1.2 mM K2HPO4, 10 mM glucose). For addition experiments, the respective compound (2-Desoxyglucose (Sigma Aldrich 25972), antimycin-A (Sigma Aldrich A8674), arachidonic acid (Sigma Aldrich 10931), ATP (Jena Bioscience NU-1049), forskolin (Sigma Aldrich F6886), ionomycin (Thermo Fisher Scientific 11415691), neurotensin ($NTS_{8-13}$ peptide synthesized by GeneScript)) was diluted in IB at 2x concentration, and 30 seconds after starting the acquisition, 50 µl were added to the cells while imaging.

Plasma membrane (PM) targeting of ras-modified GEQO2f constructs was assessed in cells by imaging single timepoints. Cells transiently expressing PM-targeted GEQO2f variants were stained using a 1:2 000 dilution of CellMask DeepRed (Thermo Fisher Scientific C10046) in IB for 5 minutes, washed 3 times with IB, and imaged at 37°C, 5%CO2 using a 60x oil immersion objective.

GEQO constructs were imaged using 0.1 % relative laser power at 445 nm and a 405-445/514 nm dichroic mirror. Emission was detected between 500-550 nm for the sensor channel and 610-700 nm for the reference channel.

Emission spectra of caGEQO, both expressed in HCT116 cells and as purified protein, were recorded using the lambda function of the Olympus FluoView software, using excitation at 445 nm at 3 % relative laser power and the 405-445/514 nm dichroic mirror. Emission was detected in 10 nm steps ranging from 490-710 nm.

**Zebrafish handling, transgenesis, and live imaging**

All zebrafish (*Danio rerio*) lines were maintained in a recirculating system with a 14 h/day, 10 h/night cycle at 28°C. Crosses were performed with 3- to 12-month-old adults. Embryos were kept in E3 zebrafish embryo medium (5 mM NaCl, 0.17 mM KCl, 0.33 mM CaCl2, 0.33 mM MgSO4, 5% Methylene Blue, pH 7.2) at 28.5°C until the desired developmental stage was reached.

For transgenesis, one-cell stage wildtype AB strain zebrafish embryos were injected using standard procedures with 25 pg of each of the GECO plasmids (see above), together with 25 pg of Tol2 transposase mRNA. The injected embryos (F0) were screened at 24 hours post fertilisation (hpf) for green and red fluorescence signals using an Olympus SZX16 fluorescence stereoscope. Positive embryos with mosaic expression were selected and grown to adulthood. To screen for positive founders (F0), adult fish were crossed with wildtype AB, and respective F1 progeny was screened for fluorescence as before. Identified F1 positive embryos were selected and raised to adulthood. The produced transgenic alleles are as follows: Tg(ubi:GEQO2-1f)cbg25tg
Tol2 mRNA was generated by linearization of pCS2FA-transposase plasmid[3] with NotI (NEB) and transcribed using the SP6 mMESSAGE mMACHINE High Yield Capped RNA Transcription Kit (#AM1340, Ambion), following manufacturer's protocol. mRNAs were aliquoted and stored at -70°C until use. A PV-820 Pico-injector (World Precision Instruments) and a Narashige micromanipulator were used for microinjection.

For confocal live imaging, 24 hpf embryos were dechorionated and anesthetized with 0.01% MS-222 (Sigma E10521) diluted in E3 medium, then mounted in 0.5% low melting point agarose (Sigma A9414) diluted in E3 fish medium, using 35mm glass bottom dishes (MatTek), placing the tail or trunk regions as close as possible to the surface of the coverslip. Live embryos were then imaged on an Olympus FluoView3000 point scanning confocal microscope at 28.5°C using a 60x water immersion objective. Images of a single z-plane were acquired for 10 min at 1 fps.

**Image analysis**

OIR files were converted to TIFF files using ImageJ. All further data processing was done using Python. All scripts can be found in the corresponding repository (https://doi.org/10.17617/3.9IHKTD).

Cells expressing GEQO2 or similar constructs were segmented based on the reference channel using Cellpose3[4,5]. To account for cell movement, the first and last frame of movies were segmented separately and non-overlapping regions were removed.

Zebrafish epidermis cells expressing GEQO2f were segmented using Cellpose3 based on a sum projection of the sensor channel over all time points.

The mean intensity of every labelled region was measured for all channels and bleach corrected based on untreated control movies recorded at identical microscope settings.

Cells expressing PM targeted versions of GEQO2f were segmented based on the plasma membrane co-stain using a local minima-seeded watershed algorithm (napari pyclesperanto-prototype). The resulting labels were filtered to only include positively transfected cells. Using an edge detection algorithm, the labels were separated into a membrane and a cytosolic region.

**Microscope calibration**

To obtain calibration curves for the conversion of fluorescence intensities to calcium and protein concentrations, eight calcium concentrations (0-3 µM) at five different protein concentrations (0-2 µM) were imaged at various imaging conditions, matching those used to image cells and zebrafish. The calcium dilutions were prepared using the Invitrogen Calcium Calibration Kit #1 as indicated above. Protein solutions were prepared as 10x concentrates and combined 1:9 with the calcium dilutions. These measurements yield three calibration curves that are then used to convert observed fluorescence intensities to absolute GEQO expression levels and analyte concentrations as follows.

**Analyte quantification**

Fluorescence intensities of GEQO sensors were converted to absolute concentrations of sensor proteins and calcium as illustrated in Extended Data Figure 2:

The bleach corrected fluorescence intensity of the reference channel was used to calculate the GEQO expression level for every cell using calibration curve A, relating reference channel fluorescence $F^{Reference}$ to the concentration of the respective GEQO version using the following equation (Extended Data 2, 1st row):

$$[GEQO] = \frac{F^{Reference} - b_A}{m_A} \quad (1)$$

The relation of the observed sensor fluorescence $F^{\text{Sensor}}$ to the GEQO and calcium concentrations, was used to estimate the maximum $F^{\text{Sensor}}$ at any given sensor concentration, with $F_{\text{max}}^{\text{Sensor}}$ being the maximum sensor fluorescence at calcium-saturated ($\text{Ca}^{2+}\text{sat}$) conditions (Extended Data 2, 2nd row):

$$F_{\text{max}}^{\text{Sensor}} = [\text{GEQO}] * m_{\text{Ca}^{2+}\text{sat}} + b_{\text{Ca}^{2+}\text{sat}} \quad (2)$$

With the GEQO expression level and $F_{\text{max}}^{\text{Sensor}}$ all required information is available to convert the observed sensor fluorescence to calcium concentrations by applying the law of mass action. Because all GEQO2 constructs bind calcium cooperatively, as indicated by the Hill coefficients of their respective dose-response curves (GEQO2f: 1.6, GEQO2m: 2.7, GEQO2s: 2.1; compare Extended Data 1g), the Hill coefficient needs to be taken into account for the conversion. To implement this dependency, the Hill-Langmuir equation was used to convert $F^{\text{Sensor}}$ to absolute calcium concentrations. The equation is frequently used to describe the cooperative binding of ligands to a macromolecule or receptor, such as the binding of oxygen to haemoglobin, as the fraction $\Theta$ of ligand-bound receptor to the free concentration of ligand:

$$\begin{aligned} \Theta &= \frac{[L]^{nH}}{K_\mathrm{d} + [L]^{nH}} \\ &= \frac{[L]^{nH}}{(K_\mathrm{A}^{nH} + [L]^{nH})} \\ &= \frac{1}{1 + \left(\frac{K_\mathrm{A}}{[L]}\right)^{nH}} \end{aligned} \tag{3}$$

Where $\Theta$ is the fraction of ligand bound receptor ($\Theta = \mathrm{Receptor_{bound}}/\mathrm{Receptor_{total}}$), $[L]$ is the concentration of free ligand, $K_\mathrm{d}$ is the apparent dissociation constant, $K_\mathrm{A}$ is the ligand concentration at half saturation, and $nH$ is the Hill coefficient.

When converting fluorescence intensities, $F_\mathrm{measured}^\mathrm{Sensor}$ indicates the concentration of calcium-bound GEQO and $F_\mathrm{max}^\mathrm{Sensor}$ corresponds to the total concentration of GEQO, making $\Theta = F_\mathrm{measured}^\mathrm{Sensor} / F_\mathrm{max}^\mathrm{Sensor}$ (Extended Data 2, 2nd row). Thus, the calcium concentration at any given timepoint was calculated (Extended Data 2, 3rd row):

$$\begin{aligned} [\mathrm{Ca}^{2+}] &= \frac{K_\mathrm{A}}{\sqrt[nH]{\frac{1}{\Theta} - 1}} \\ &= \frac{K_A}{\sqrt[nH]{\frac{1}{\left(\frac{F_{measured}^{Sensor}}{F_{max}^{Sensor}}\right)} - 1}} \end{aligned} \tag{4}$$

**Direct comparison of relative and absolute quantification**

To directly assess the impact of the quantification method used to analyse data, calcium response movies of developing zebrafish epidermis cells, ATP and arachidonic acid-treated ePE cells were analysed as normalised intensiometric or ratiometric data or converted to absolute sensor and calcium concentrations. In brief: The bleach corrected fluorescence intensities of the sensor channel were either normalised to initial fluorescence yielding $\Delta F/F_0$ values for each timepoint, divided by the reference channel fluorescence intensity of the according time point and normalized to the initial fluorescence ratio yielding $\Delta R/R_0$, or converted to absolute analyte concentrations using *in-vitro* calibration curves, pre-recorded at identical microscope settings. The resulting data sets were then analysed using the parametrisation routine described below. Thus, it was possible to directly compare intensiometric, ratiometric, and absolute quantified data.

**Response parametrisation**

Calcium responses of individual cells were described using an automatically generated feature set extracted from absolute quantified, ratiometric, and intensiometric data sets. First, responses were identified using thresholding. The threshold value was calculated for every cell individually, based on the standard deviation of the mean and a manually set scaling factor to random noise in the data (scaling factors for different experiments: HCT 116 cells: 7.5, ePE cells: 7.5, *D. rerio*: 3). For experiments using compound addition to induce calcium responses the baseline was calculated based on all time points before compound addition. For *D. rerio* experiments the baseline was calculated by excluding the top 10% of calcium concentrations. For every response, the maximal calcium concentration (or $\Delta F/F_0$ or $\Delta R/R_0$ value) was determined as the peak amplitude and the response duration was determined as the time from response onset (first timepoint surpassing the threshold) to response end (last timepoint surpassing the threshold). The time elapsed from response onset to response peak was defined as the rise time. The time elapsed from response peak to response end was defined as the decay time. The time elapsed between compound addition and response onset was defined as the lag time. The calcium concentration at resting state of every cell was determined as the mean value before induction of calcium responses. As additional parameters, calcium traces were described by the area under the curve of the baseline subtracted calcium concentration (or $\Delta F/F_0$ or $\Delta R/R_0$ value). The area under the curve was calculated for individual responses and globally for the total time trace.

**Response analysis**

The parametrised calcium responses were further analysed using different dimensionality reduction and clustering approaches. Uniform Manifold Approximation and Projection (UMAP) dimension reduction was applied as implemented in the umap package for Python [6]. Principal component analysis (PCA) was applied as implemented in the Python package scikit-learn.decomposition.PCA. The number of components for PCA was chosen to achieve an explained variance >70%. K-means clustering was applied as implemented in the Python package scikti-learn.cluster.KMeans. The number of clusters was chosen to minimise the within cluster sum of squares while maximising the silhouette score calculated using scikit-learn.metrics.silhuette_score.

**Signal-to-noise and bleed-through analysis in cells**

To evaluate the signal-to-noise ratio (SNR) and bleed-through in cells, calcium responses were elicited by sequentially adding neurotensin (NTS) to trigger physiological calcium release from internal stores, followed by ionomycin to fully saturate the sensor response.

The baseline fluorescence intensity in the sensor channel was defined as the mean fluorescence value over the first 25 timepoints (prior to stimulation). The SNR was calculated as the ratio of the maximum fluorescence intensity recorded in the sensor channel to the standard deviation of the baseline, following the methodology described by Akerboom et al. (2012)[7].

Bleed-through, defined as fluorescence emitted by the sensor domain but detected in the reference channel, was quantified by comparing the fluorescence signals in the two channels. To account for the entire imaging period, the area under the curve (AUC) of fluorescence intensity over time was calculated for each channel. The ratio of the cumulative fluorescence intensity (AUC) in the reference channel to that in the sensor channel was used as a measure of bleed-through.

**Plasma membrane targeting analysis**

The efficiency of plasma membrane targeting of different plasma membrane targeting domains was analysed for single cells using two complementary methods.

The ratio of the mean fluorescence intensity (normalised by area) of the plasma membrane label and the cytosolic label was calculated as $PM/Cytosolic\ Ratio\ =\ F^{PM}/F^{Cytosol}$.

For each cell, the Pearson correlation coefficient between the GEQO sensor channel and the CellMask DeepRed channel was calculated using the python package bebi103 by Justin Bois[8] using the Costes colocalization method[9].

### Statistical analysis

Statistical analysis was performed using two-sided bootstrap hypothesis testing with resampling, implemented in a custom Python function that estimates p-values using a normal approximation via scipy.stats.norm. Significance levels are indicated as follows:

***: p < 0.001,

**: p < 0.01,

ns: p > 0.05
